# Classification models for Invasive Ductal Carcinoma Progression, based on gene expression data-trained supervised machine learning

**DOI:** 10.1101/666222

**Authors:** Shikha Roy, Rakesh Kumar, Vaibhav Mittal, Dinesh Gupta

## Abstract

Early detection of breast cancer and its correct stage determination are important for prognosis and rendering appropriate personalized clinical treatment to breast cancer patients. However, despite considerable efforts and progress, there is a need to identify the specific genomic factors responsible for, or accompanying Invasive Ductal Carcinoma (IDC) progression stages, which can aid the determination of the correct cancer stages. We have developed two-class machine-learning classification models to differentiate the early and late stages of invasive ductal carcinoma. The prediction models are trained with RNA-seq gene expression profiles representing different IDC stages of 610 patients, obtained from The Cancer Genome Atlas (TCGA). Different supervised learning algorithms were trained and evaluated with an enriched model learning, facilitated by different feature selection methods. We also developed a machine-learning classifier trained on the same datasets with training sets reduced data corresponding to IDC driver genes. Based on these two classifiers, we have developed a web-server Duct-BRCA-CSP to predict early stage from late stages of IDC based on input RNA-seq gene expression profiles. The analysis conducted by us also enables deeper insights into the stage-dependent molecular events accompanying breast ductal carcinoma progression. The server is publicly available at http://bioinfo.icgeb.res.in/duct-BRCA-CSP.

## Introduction

Breast cancer ranks second among all the cancer types arranged in the order of increasing death rates, also the most prevalent cancer in women [1]. The cancer has been categorized into three therapeutic groups: ER - ER+ patients receive endocrine therapy, HER – HER+ group is treated by therapeutic targeting of HER/ERBB2, and TNBC - lacking expression of ER, PR, HER receptors [2]. It has been categorized into two major histological types-Invasive Ductal Carcinoma (IDC) and Invasive Lobular Carcinoma (ILC), occurring in 47-79% and 2-15% of invasive cancers amongst women of different worldwide races, respectively [3], [4]. These two sub-types show similarity in certain features such as tumour site, tumour size, stage and grade, but have different metastatic patterns, characteristic histology and malignant calcifications [5], [4]. IDC starts from ducts and spreads to the breast fatty tissue, whereas ILC is restricted to milk producing lobules [6]. These two sub-types are also discriminated at the molecular level with differential expression of gene encoding vimentin, cathespin D, thrombospondin, E-cadherin, vascular endothelial growth factor, cytokeratin 8, and cyclin A. [4], [7], [8], [9],[10], [11], [12]. The pathological differences between the two sub-types arises as a result of separate gene regulatory networks, which warrants further exploration for the development of appropriate diagnostic and therapeutic treatment strategy [6]. According to reports, 75% cases of invasive breast carcinoma cases are accounted by IDC, however, advanced treatment of IDC patients still remains a challenge due to lack of molecular targets for IDC treatment [13], [14]. Also, there is the availability of higher number of datasets for IDC patients in TCGA-BRCA, which is favourable for development of efficient classifiers using machine learning. Hence, we implemented machine-learning and developed a web-server for efficient prediction of the correct IDC stage, which can potentially aid in designing appropriate treatment strategies and precise molecular targeting.

The increased incidence of breast cancer and higher mortality rate has attracted significant research efforts to unravel its causes, and development of better treatment options [15], [16]. Breast cancer is a heterogeneous disease with varied features, such as morphological appearances, profile, response to therapy, TNM staging, histological grade, etc. [17]. There is a direct correlation between mortality rate and stages of cancer, and the stage progression could be checked by early detection and appropriate treatment strategies [18]. Although knowledge about genomic profiling has been identified in terms of varied molecular features associated with subtypes of cancer, its molecular mechanism of progression is poorly understood [19]. Tumour stage is defined as the anatomic extent of cancer at the time of diagnosis, which is important for an individual patient prognosis, and determination of best treatment strategy [20]. Pierre Denoix and the Union of International Cancer Control (UICC) has classified tumour staging based on TNM classification [20]. TNM classification overlaps with breast cancer stages, where T describes the extent of a primary tumour by the size or depth of invasion mainly in stage I or II, N describes the extent of regional lymph node metastasis in mainly stage II or III, and M describes the presence of metastasis mainly in stage IV [20]. The incorporation of this staging system into molecular or genetic profiles can help in detecting prognostic groups that guide the disease intervention [20]. There is a sharp decrease in the 5-year survival rate of patients with the stage-wise progression of breast cancer [18]. Treatment of cancer remains a challenge because of the lack of knowledge about factors for cancer progression and metastasis [20]. Potential treatment options are available based on clinical and pathological prognostic factors with the histological grade being the most important predictive factor [20]. High throughput techniques such as Next Generation Sequencing (NGS) that capture expression of thousands of genes in a single assay can act as powerful analytical tools for capturing breast cancer prognostic signature [17]. We can obtain information about a large number of genes, but their intertwining relationship cannot be captured by traditional techniques like statistical and correlational analyses, hence advanced methods such as machine-learning are important to capture cryptic signatures inherent in these data [15], [16]. Molecular profiling helps in finding predictive information and identifying prognostic biomarkers that can serve as therapeutic targets [17]. Most of the cancer research is focussed to determine for finding driver genes, which are related to chimeras or splice junctions, which do not utilize the high-resolution features of RNA-seq [21]. Machine-learning techniques are increasingly being used for modelling the progression and treatment of cancer due to its ability to detect key features from complex datasets [22]. Personalized treatment strategies could be developed for patients with similar molecular subtypes based on the patterns identified from systematically collected molecular profiles of tumour samples [23]. In this study, we developed classification methods to analyse the genomic datasets of invasive ductal carcinoma obtained from TCGA, using supervised machine-learning algorithms and feature selection methods. We developed prediction models that could discriminate between early and late stages of IDC using RNA-seq datasets. Different feature selection methods such as RFE, RLASSO, linear modelling, linear regression and random forest were trained and evaluated using Python scikit-learn library which provides individual rankings to gene features. Based on the most comprehensive ranking of gene features by various feature selection methods the top gene features were selected for enriched classifier training that helped us efficiently classify the tumours based on the tumour stage-specific gene expression profiles.

## Results

The workflow followed in our study is shown in Figure 1. The TCGA level 3 RNA-seq datasets representing 1,093 breast cancer patients were retrieved using the TCGA2STAT R package [24]. The datasets represent 610 IDC patients, the distribution of samples across testing and training set by tumour stage is given in the Table 1. TCGA2STAT package merges the molecular profile information with clinical information into a data frame that is ready for supervised machine-learning. Each of the molecular profiles consists of RNA-seq gene expression data of 20,505 genes. The import dataset consists of ‘expression’ representing the gene expression profiles of patients in terms of RPKM values (described in methods), ‘clinical data’ which consists of clinical information related to patients, and ‘merged data’ in which both the information is mapped. Samples without clinical stage assignments were excluded from our study. Samples bearing clinical stages of stage I and II were pooled together as ‘early stage’, while the stages III and IV were pooled together as ‘late stage’. We generated gene expression data frames as comma separated value (CSV) format from the data retrieved using TCGA2STAT R package, with 20505 genes as column labels and 610 TCGA patient IDs as row labels. The values obtained by mapping the reads to genome generated as gene expression estimates were used as feature vectors for training the machine-learning classifier. Hence, the entire dataset consists of a gene expression data frame with a dimension of 610 * 20505. Near zero variance features and features having correlation coefficient more than 80% were removed using caret, an R package [25]. This led to a preliminary reduction of the number of features from 20,505 to 17,373. The training datasets were standardized using z-score normalization. It converts all the features to common scale with mean zero and standard deviation 1 (Figure S1,S2 in the Supplementary file I). The normalized data-set was used for model generation to discriminate early versus late stages of the cancer.

**Figure 1:**
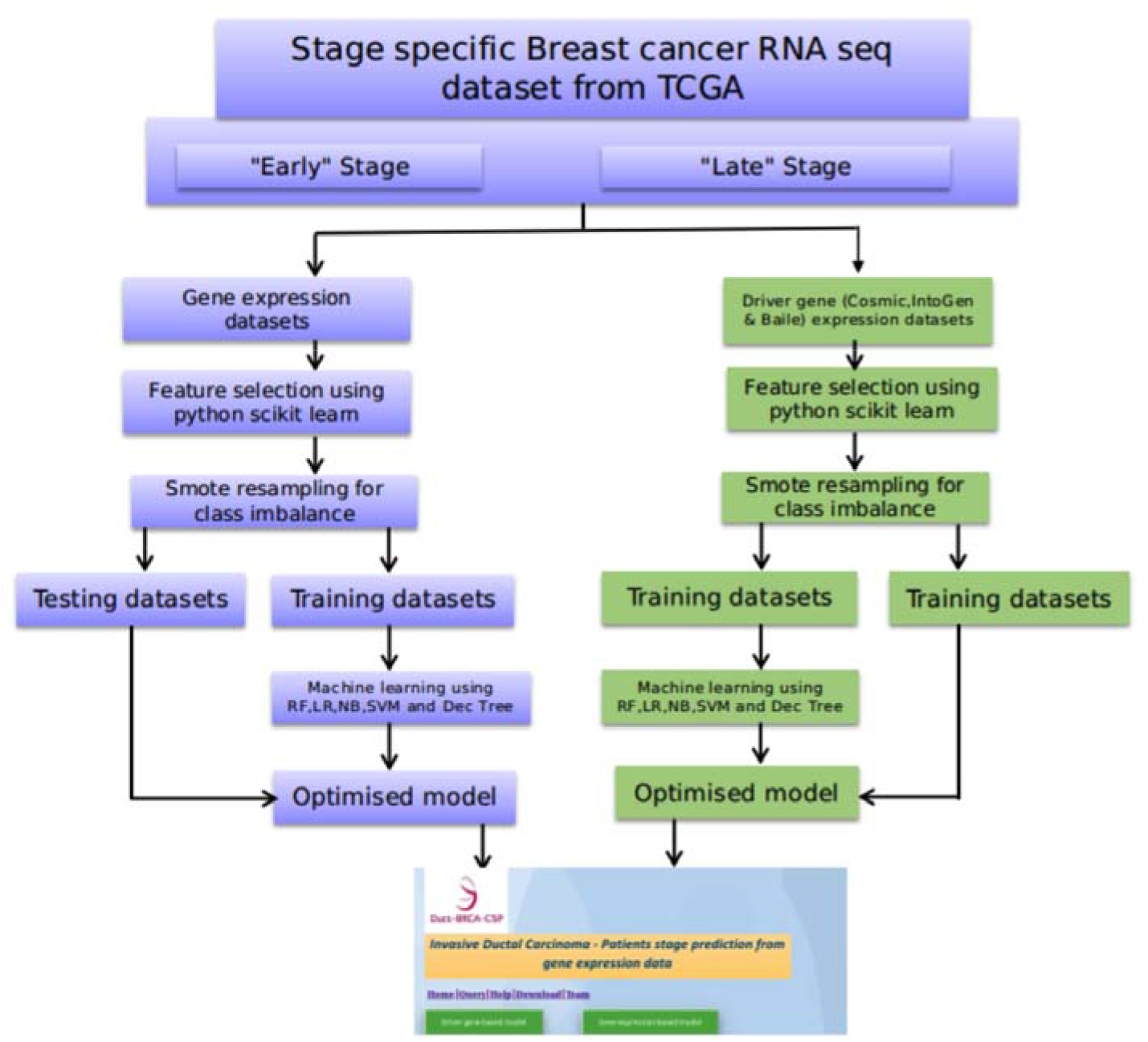
Duct-BRCA-CSP development pipeline. Study flowchart for development of classification models, trained with relevant gene expression profiles to efficiently discriminate between the early and late IDC stages.

**Table 1.**
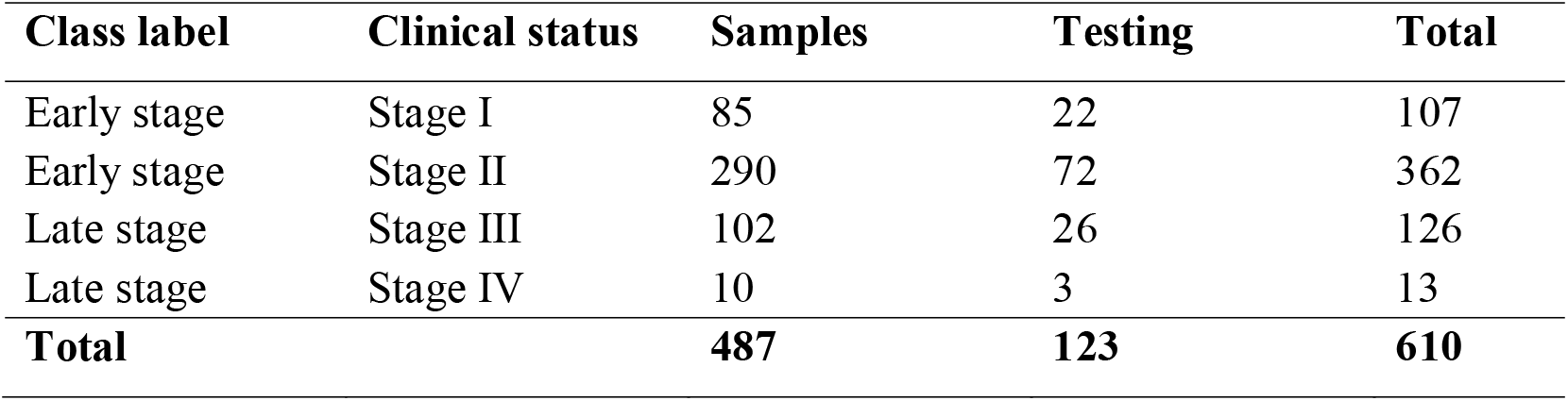
Summary of the training and testing data-sets for each stage.

The normalized datasets were divided into two training sets, the first dataset comprises of complete gene expression datasets which were the original datasets representing expression of 17,373 genes used for feature selection. The second dataset consists of gene expression data corresponding to the driver-gene list in which the training genes were reduced to driver-genes responsible for progression of different cancers. The list of 881 driver genes was obtained from three well curated driver genes lists- Cosmic, IntoGen and Bailey [26-28]. The gene expression of the selected genes of the two datasets were further used for feature selection and classifier model generation (for details, see methods section).

The top 30 gene feature list enriched models rendered the highest accuracy for driver gene expression for all the machine-learning methods, hence, these features were used for training the model (Figure 2a; 2b). The relevance of selected gene features was further validated by survival Kaplan-Meir estimate. Survival estimate revealed that median survival in cases with alteration 95.63 months and cases without alteration 129.6 months (Figure S7, Supplementary file II). Top 20 gene feature enriched models gave the highest accuracy for the complete gene expression-based model for all the machine-learning methods hence, these features were used for training the models (Figure 2c; 2d). Survival estimate revealed that median survival in cases with alteration months 128.98 months and cases without alteration 129.6 months (Figure S8, Supplementary file II). We also performed a literature validation of the selected gene features to assess the role of the selected genes in cancer progression (Supplementary file III). Despite using relevant features important for efficient training, the accuracy was low as the dataset was not balanced, i.e., there are more samples representing early stage as compared to that of late stage (469 for early stage, 141 for late stage). In order to tackle the class imbalance in the dataset, we employed Synthetic Minority Oversampling Technique (SMOTE) using Python scikit-learn library. SMOTE was employed using ENN (Edited Nearest Neighbour) in which oversampling and under-sampling is performed until there is no difference with k-neighbour of majority class [29]. Real world datasets have higher composition of ‘normal class’ as compared to ‘abnormal class’, introducing bias in classification model. Combination of over-sampling of minority class along with undersampling of majority class can aid in increasing the classifier performance [30]. To check the SMOTE resampling, models were trained on datasets where SMOTE resampling was employed (Figure S5 in the Supplementary file I). The dataset where SMOTE was employed, the classification accuracy improved from 77% to nearly 89% on the validation set (Figure S6 in the Supplementary file I). For training, validation, and testing, the samples were randomly stratified and split into 80% training-cum-validation sets (datasets available on the duct-BRCA-CSP webserver) and 20% independent testing datasets (available on duct-BRCA-CSP webserver).

**Figure 2:**
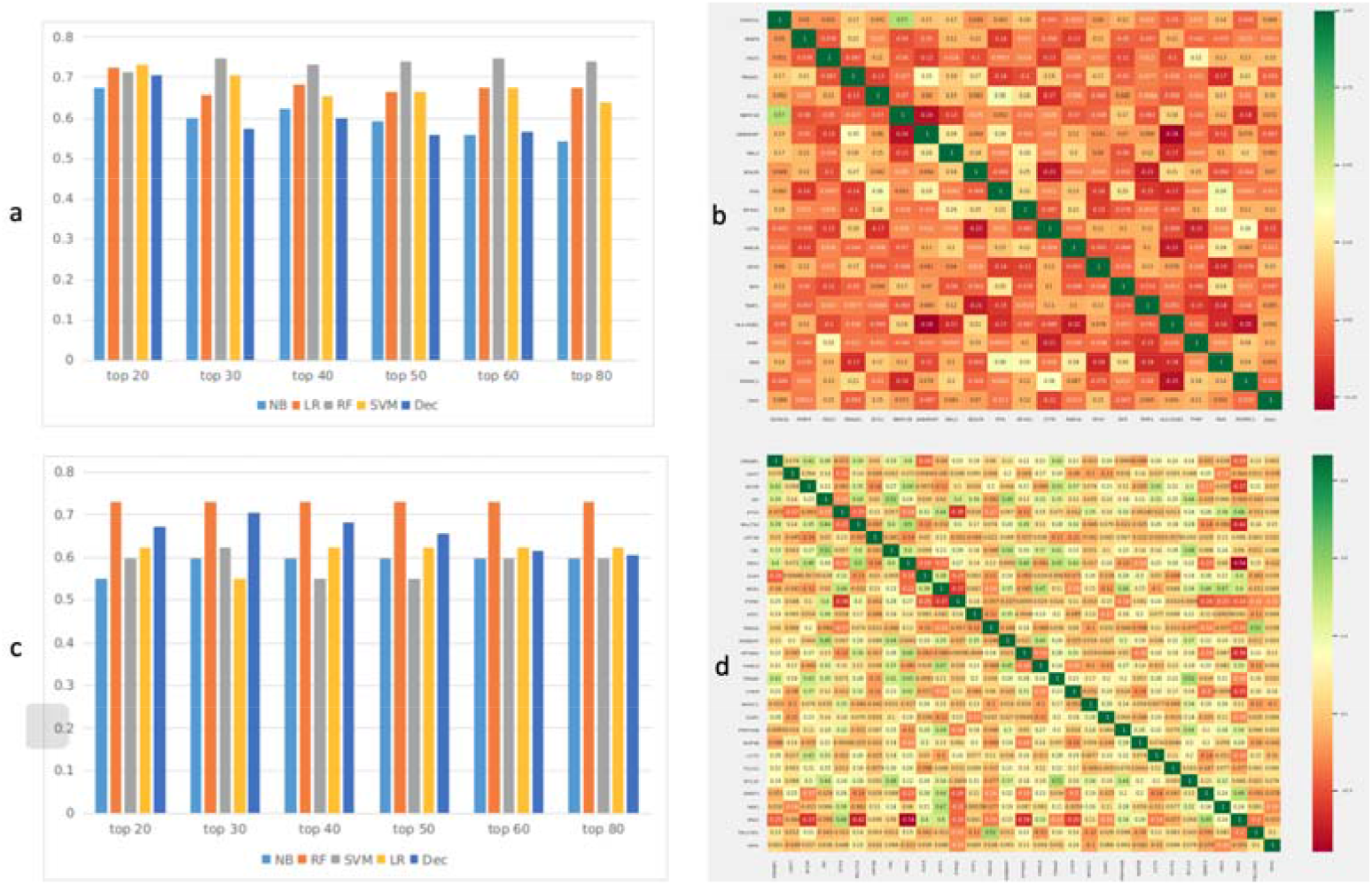
Feature selection for training datasets. 2: a. Feature selection methods were used to rank the gene features used in the training datasets. Top 50, 60, 80 and 100 features were used to train the binary classification model and their accuracy was evaluated. Based on that, top 20 gene features render highest accuracy for all the machine learning algorithms evaluated by us. NB: Naïve Bayes, LR: Logistic Regression, RF: Random Forest, SVM: Support Vector Machine, DT: Decision Tree. X-axis: model accuracy, Y-axis: no. of features selected for model building b. Correlation plot for top 20 gene features used in classification model building. X axis: Genes selected by feature selection Y axis: Genes selected by feature selection. c. Feature selection methods were used to rank the gene features used in the training datasets. Top 50, 60, 80 and 100 features were used to train the binary classification model and their accuracy was evaluated. Based on that, top 30 gene features renders highest accuracy for all machine learning algorithms. NB: Naïve Bayes, LR: Logistic Regression, RF: Random Forest, SVM: Support Vector Machine, DT: Decision Tree. X-axis: model accuracy Y-axis: no. of features selected for model building d. Correlation plot for top 30 driver gene features used in model building X, Y axis: Genes selected by feature selection.

### Training-cum-validation

The classification accuracy of the generated prediction models ranges from 74% for SVM, to 95% for Random Forest; and auROC value ranges from 0.76 for LR to 0.93 for the Random Forest trained model for complete gene expression-based model. Based on the model accuracy and auROC, we inferred that the Random Forest based prediction model has outperformed the other four machine-learning algorithms implemented in the study (Table 2). Random forest based model achieved the best performance with ROC of 0.93 on the training dataset, evaluated using ten-fold cross-validation for the complete gene expression-based model (Figure 3a). The Random forest model displayed highest auROC as compared to the other models for complete gene expression-based model (Figure 3b). The classification accuracy of the generated prediction models ranges from 72% for SVM, to 92% for Random forest; and auROC value ranges from 0.72 for LR to 0.96 for Random forest for driver gene expression-based model. Based on accuracy and auROC, we inferred that Random forest based prediction model has outperformed the four other machine-learning algorithms implemented in the study. (Table 2). Random forest based model achieved maximum performance with R0C of 0.96 on training dataset when evaluated using ten-fold cross-validation for driver gene expression-based model (Figure 4a). Random forest model exhibited the highest area under the curve as compared to the other models for driver gene expression-based model (Figure 4b).

**Table 2:** Performance of prediction model generated by ten-fold cross validation on training cum validation datasets

| Training set | Model | ACC | SEN | SPC | MCC | auROC |
| --- | --- | --- | --- | --- | --- | --- |
| Selected Gene expression | RF | 95 | 96 | 92 | 0.86 | 0.93 |
|  | DT | 85 | 86 | 79 | 0.61 | 0.77 |
|  | NB | 75 | 79 | 60 | 0.35 | 0.77 |
|  | LR | 74 | 74 | 66 | 0.23 | 0.76 |
|  | SVM | 74 | 73 | 95 | 0.29 | 0.80 |
| Driver gene expression | RF | 92 | 90 | 98 | 0.82 | 0.96 |
|  | DT | 82 | 84 | 77 | 0.60 | 0.80 |
|  | NB | 72 | 75 | 63 | 0.34 | 0.75 |
|  | LR | 73 | 74 | 66 | 0.35 | 0.72 |
|  | SVM | 76 | 76 | 76 | 0.43 | 0.76 |

Accuracy (ACC), sensitivity (SEN) and specificity (SPC) values in %

**Figure 3:**
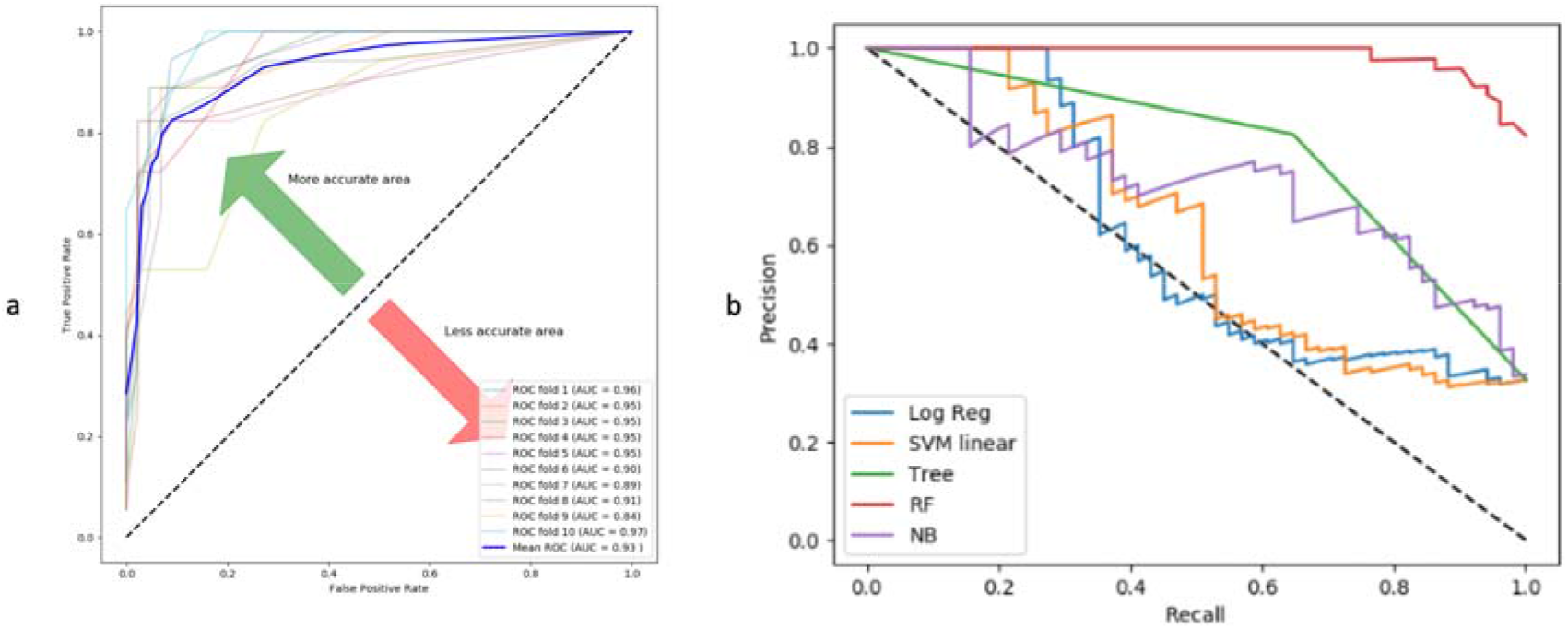
ROC and Precision-recall curve for gene-expression based models. a. Random forest based model achieved maximum performance with R0C of 0.93 on training dataset when evaluated using ten-fold cross-validation for complete gene expression-based model Receivers Operating Curve (ROC) of the Random forest classifier with 10-fold cross validation which is having highest accuracy. b. Precision-recall curve is a trade of between precision and recall with high area under the curve representing low false positive and low false negative for all classifiers. Amongst all the prediction models, Random Forest achieved the maximum area under precision-recall curve for complete-gene expression model.

**Figure 4:**
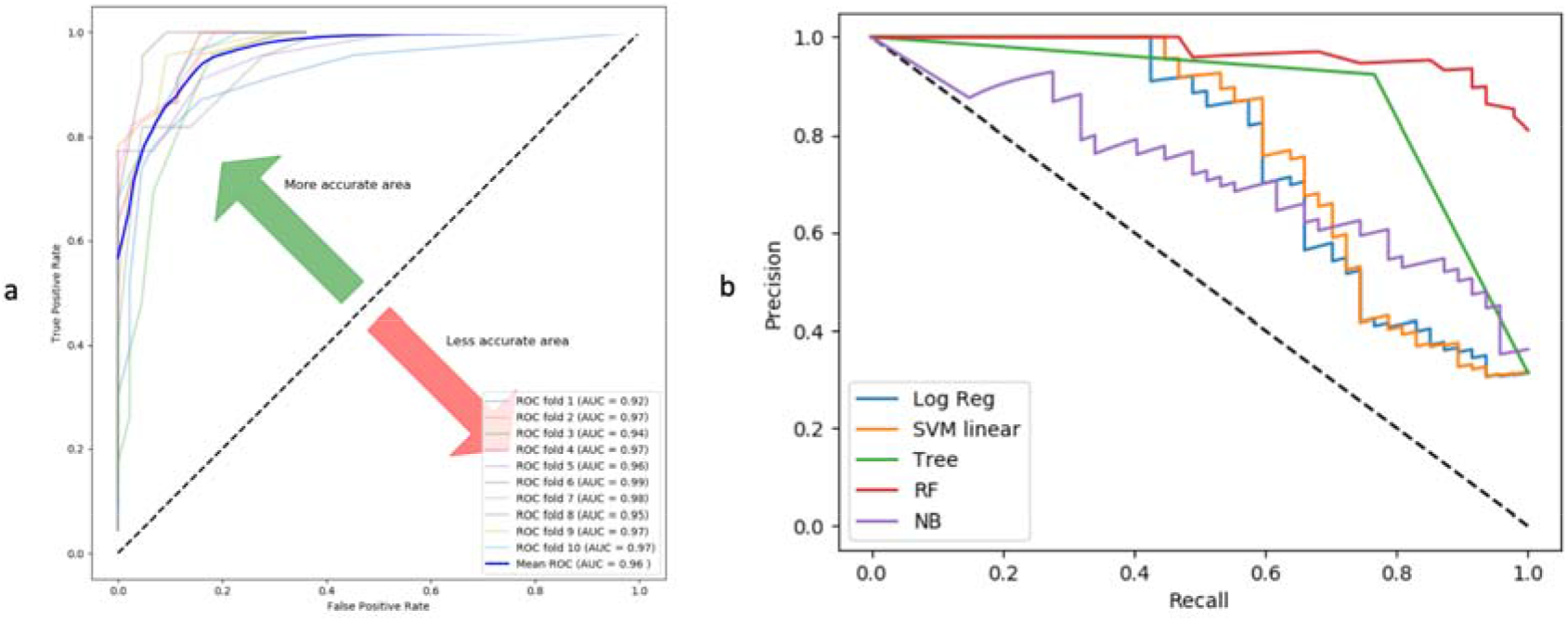
ROC and Precision-recall curve for driver gene expression-based model. a. Random forest based model achieved maximum performance with R0C of 0.96 on training dataset when evaluated using ten-fold cross-validation for driver gene expression-based model Receivers Operating Curve (ROC) of the Random forest classifier with 10-fold cross validation which is having highest accuracy. b. Precision-recall curve is a trade of between precision and recall with high area under the curve representing low false positive and low false negative for all classifiers. Amongst all the prediction models, Random Forest achieved the maximum area under precision-recall curve for driver-gene expression-model.

### Independent data-set performance

Further, we evaluated the performance of the trained models on independent datasets. The performance was re-evaluated based on accuracy, sensitivity, specificity, MCC and auROC for all the models. We observed coherence in the performance of the models between independent data testing and 10-fold cross validation based on auROC values for the complete gene expression-based model. Random forest achieved maximum ROC of 0.969 with an accuracy of 90% for testing datasets implemented in the complete gene expression-based model (Table 3). Also, we observed coherence in the performance of the models between independent data testing and 10-fold cross validation based on auROC values for driver gene expression-based model. Random forest achieved maximum auROC of 0.99 with an accuracy of 94% for testing datasets in driver gene expression-based model (Table 3).

**Table 3:** Performance of prediction models by standard statistical evaluation parameters for independent testing dataset

| Training set | Model | ACC | SEN | SPC | MCC | auROC |
| --- | --- | --- | --- | --- | --- | --- |
| Selected Gene expression | RF | 90 | 89 | 91 | 0.78 | 0.96 |
|  | DT | 84 | 84 | 84 | 0.65 | 0.81 |
|  | NB | 71 | 73 | 65 | 0.35 | 0.82 |
|  | LR | 67 | 67 | 71 | 0.23 | 0.62 |
|  | SVM | 74 | 72 | 1 | 0.36 | 0.57 |
| Driver gene expression | RF | 94 | 94 | 1 | 0.88 | 0.99 |
|  | DT | 84 | 84 | 83 | 0.65 | 0.81 |
|  | NB | 73 | 75 | 71 | 0.42 | 0.82 |
|  | LR | 74 | 74 | 76 | 0.43 | 0.75 |
|  | SVM | 77 | 75 | 87 | 0.51 | 0.73 |

Accuracy (ACC), sensitivity (SEN) and specificity (SPC) values in %

### External validation for a microarray dataset

We also evaluated the performance of the models developed by us for another dataset representing a microarray data, obtained from GEO. The models were able to achieve a maximum auROC of 0.46 with an accuracy of 67% for the Random forest based model (Table 4). A maximum ROC of 0.45 with accuracy 38% with Random forest based model trained on driver gene expression features (Table 4). Heatmap of differential expression analysis of microarray datasets between early and late stage for the complete gene expression-based features set (Figure S3, Supplementary file I); and driver gene-based features set, showing differences in gene expression between early and late stages for the selected gene features (Figure S4, Supplementary file I).

**Table 4:**
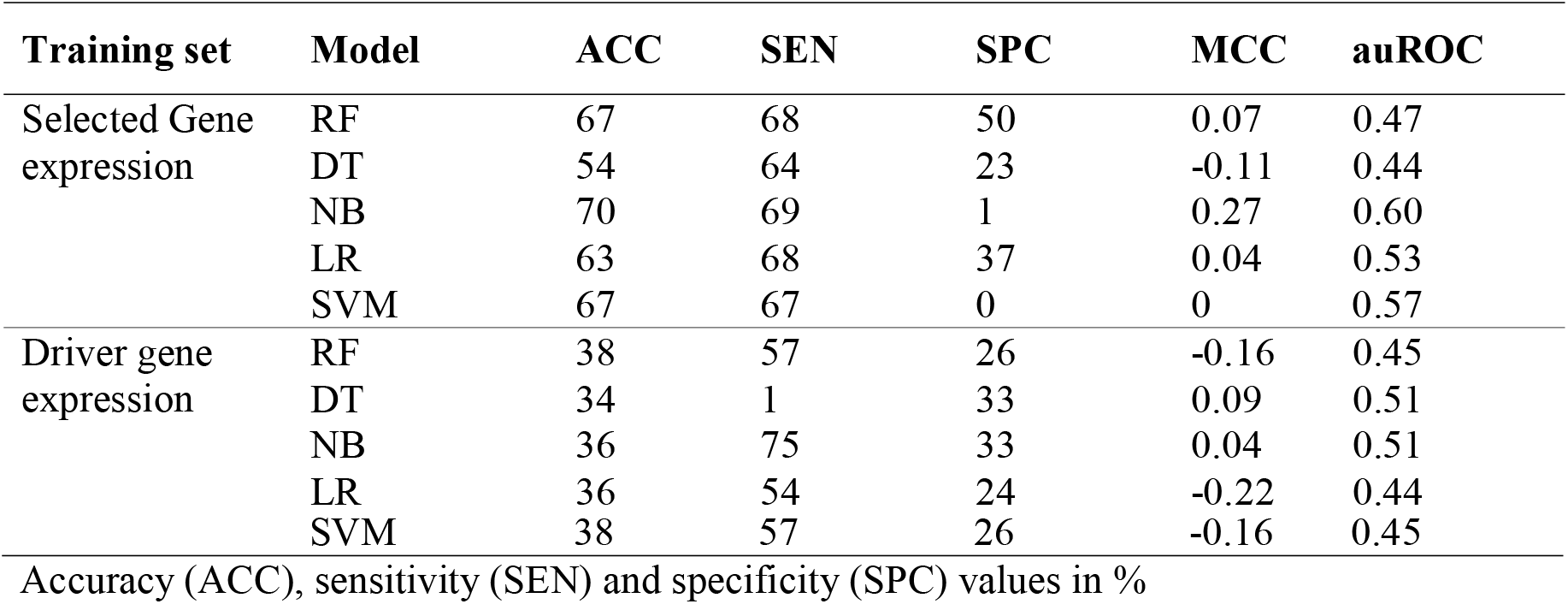
Performance of prediction models by standard statistical evaluation parameters for external validation dataset

### t-SNE (T-distributed Stochastic Neighbour Embedding)

t-SNE technique was used for visualization of our gene expression datasets that displays high-dimensional data providing each data point a location in 2D or 3D space It helps to model features into high-dimensional object to three-dimensional space such that similar objects tend to cluster together and dissimilar ones are modelled to distant points. The t-SNE analysis on our datasets segregates samples representing early and late stages, which shows that the dataset features are separable (Figure 5).

**Figure 5:**
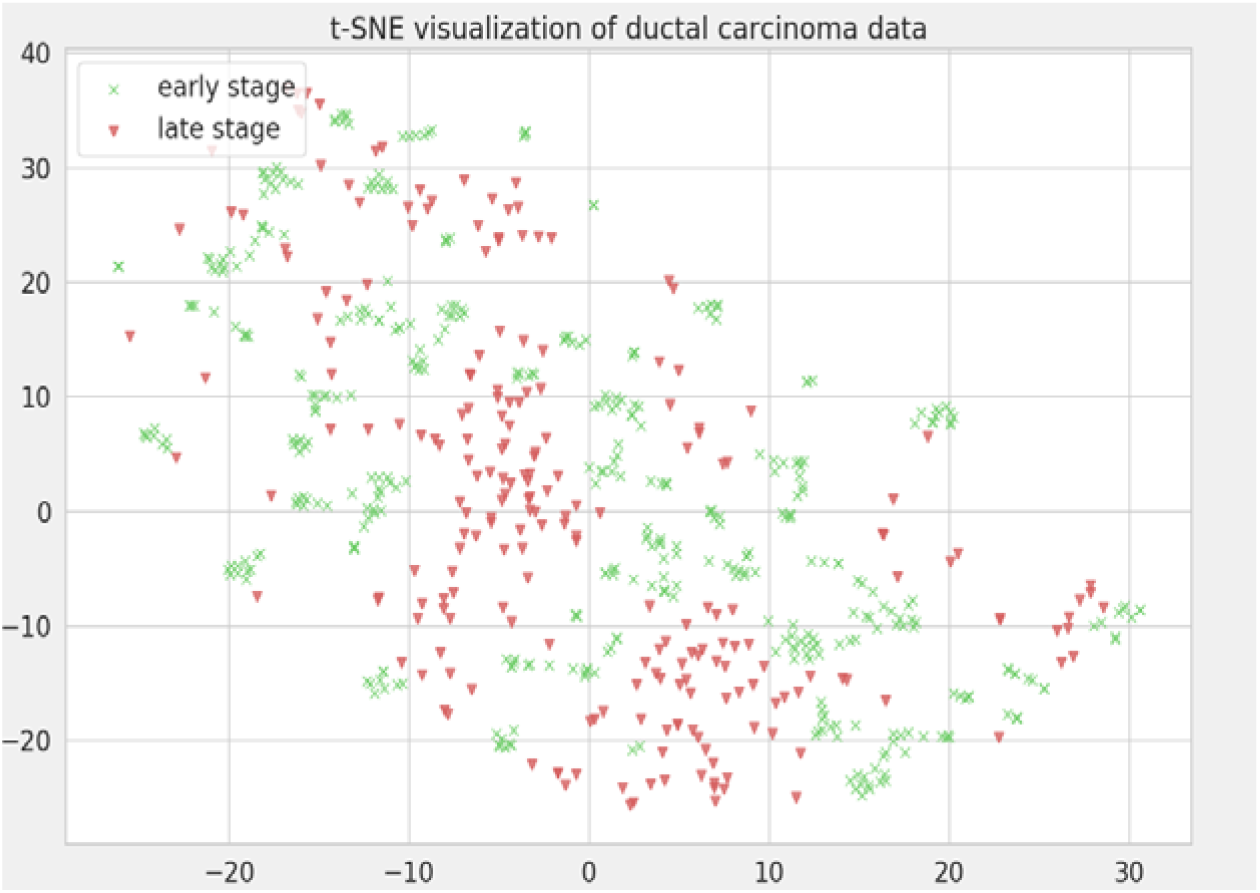
t-SNE visualization of gene expression data. t-SNE visualization was implemented on our gene expression data-sets to check if data-sets are segregating to early stage and late stage class labels based on selected features. This technique visualizes our data-sets in 3D space in which early stage and late stage samples are segregating. X axis: X in t-SNE Y axis: Y in t-SNE

### Protein-protein interaction analysis of genes selected for model building

We performed protein-protein interaction analysis on gene features selected by our models using STRING database (Search Tool for Retrieval of Interacting Genes): the complete gene expression-based model, driver gene-based model and the combination of two. We found that as compared to the former two gene sets, more interacting partners are exhibited by string analysis of their combination (Figure 6a-c). Thus, we were able to decipher major pathway that were targeted by gene sets in IDC selected by our models.

**Figure 6:**
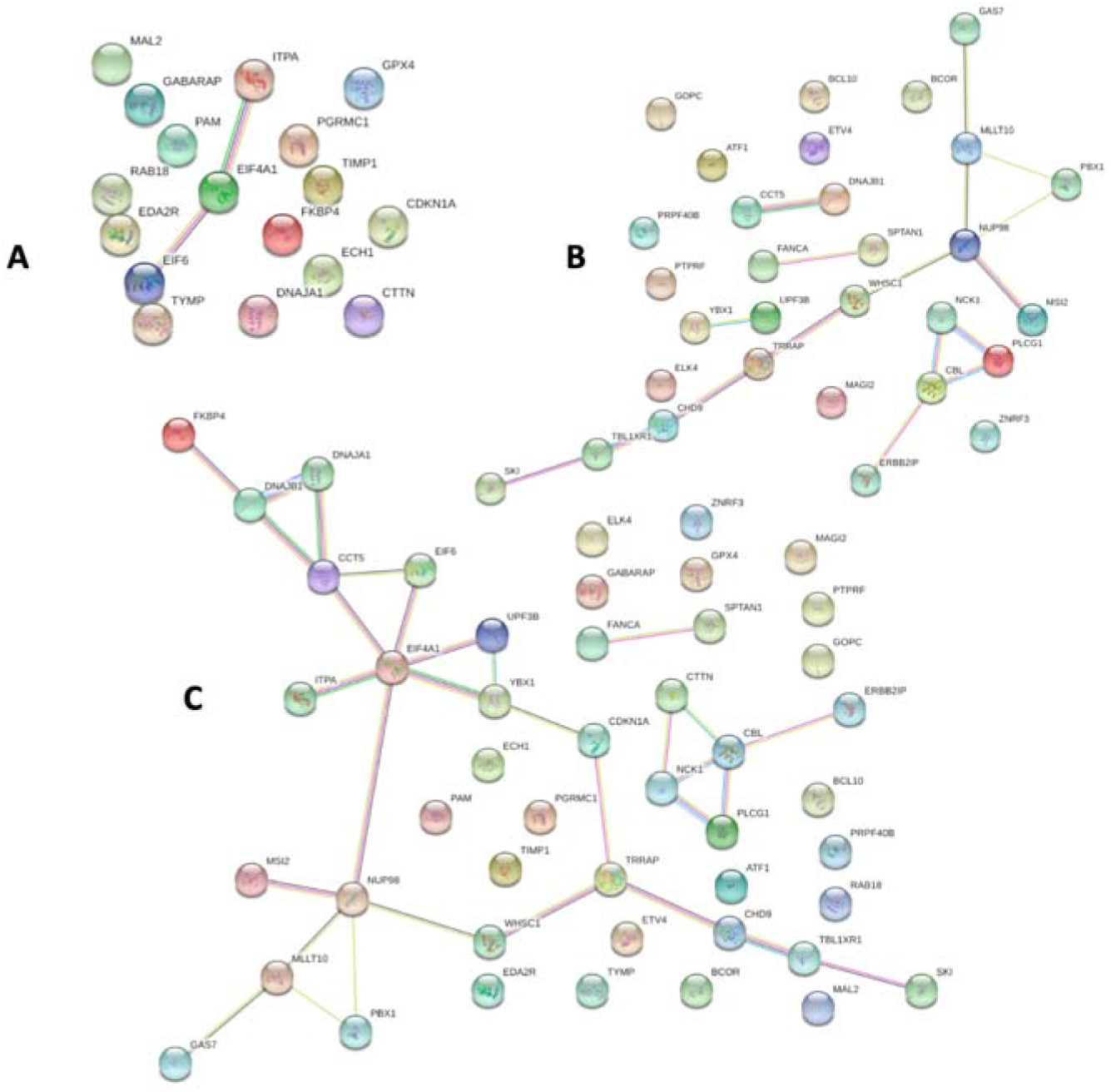
Protein-protein interaction analysis using STRING. a. Protein-protein interaction analysis using STRING of the gene set for the complete gene expression based-model. b. STRING protein-protein interaction analysis of gene set from driver gene expression-based model. c. STRING protein-protein interaction analysis of the combined gene sets. We found that as compared to the 11a and 11b, more interacting partners are exhibited by string analysis of their combination 11c which helps to decipher major pathways associated with IDC progression.

Four proteins encoded by DNAJB1, DNAJA1, CCT5 and FKBP4 are revealed to be in direct interactions, using STRING analysis. These proteins are major components of ubiquitin protein conjugation pathway by interacting with heat shock protein (Figure 6c). This process mediate cellular processes such as protein localization, cell cycle regulation and DNA damage repair [31]. Ubiquitin dys-regulation can affect tumour suppressor or oncogene leading to cellular transformation and cancer [32]. DNAJB1 binds to mitogen-inducible gene MIG6, a tumour suppressor, which positively regulates epidermal growth factor signalling, leading to breast cancer development [33].

Five proteins encoded by CTTN, NCK1, CBL, PLCG1 and ERBB2IP depicts direct interaction in STRING analysis involved in RTK signalling pathway (Figure 6c). Its aberrant expression results in enhanced cell proliferation, survival and metastasis leading to malignancy [34]. CTTN encodes cortactin which is a substrate for tyrosine Src nonreceptor tyrosine kinase whose amplification has been reported in primary metastatic breast carcinoma [35].

Four proteins encoded by TRAAP, CDKN1A, CHD9 and WHSC1 depict direct interaction in string analysis involved DNA replication and DNA damage repair pathway (Figure 6c). TRAP bind to proliferating cell nuclear antigen (PCNA) resulting DNA replication inhibition and cell growth inhibition and cancer [36]. WHSC1 is a methyl transferase that performs histone methylation affecting cell ability to undergo DNA damage repair [37].

Five proteins encoded by EIF6, ITPA, YBX1, UPF3B and EIF4A1 depict direct interaction in string analysis involved in protein translational machinery (Figure 6c). Deregulated protein synthesis can affect several processes such as cell growth, proliferation, apoptosis at translational level and malignancy [38]. Dys-regulation of EIF4A1 protein results in preferred translation of gene involved in pro-oncogenic signalling [39].

Proteins encoded by GAS7, NUP98, MSI2, MLLT10 and PBX1 depict direct interaction in string analysis involved dys-regulated DNA binding transcription factor pathway (Figure 6c). DNA binding TFs are commonly deregulated in cancer which modulates gene expression resulting IN malignancy [40]. MSI2 directly regulates estrogen receptor by binding to ESR1 resulting in breast cancer cell growth [41].

### Threshold value of expression for genes selected by feature selection

Threshold value is the expression value beyond which the sample will segregate into two groups, in our study- ‘early’ and ‘late’ stages. For example, if Z-score of CDKN1A (overexpressed in early stage) is greater than 0.32 is then it is representative of an early stage sample otherwise if it is less than 0.32 then it is representative of a late stage sample. We calculated threshold for all the genes selected by feature selection methods for the complete gene expression-based model as well as driver gene-based model (Table 5).

**Table 5:**
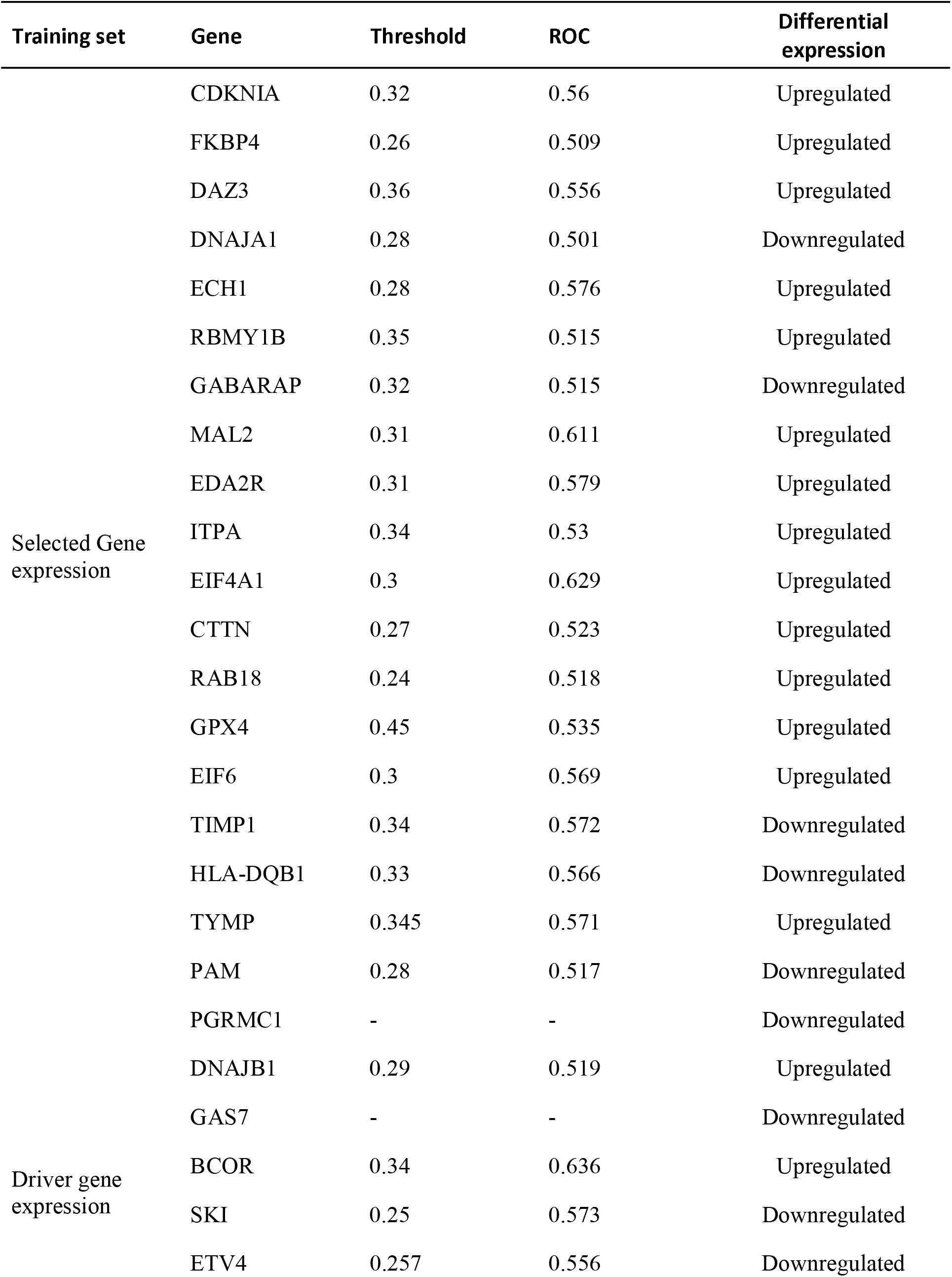
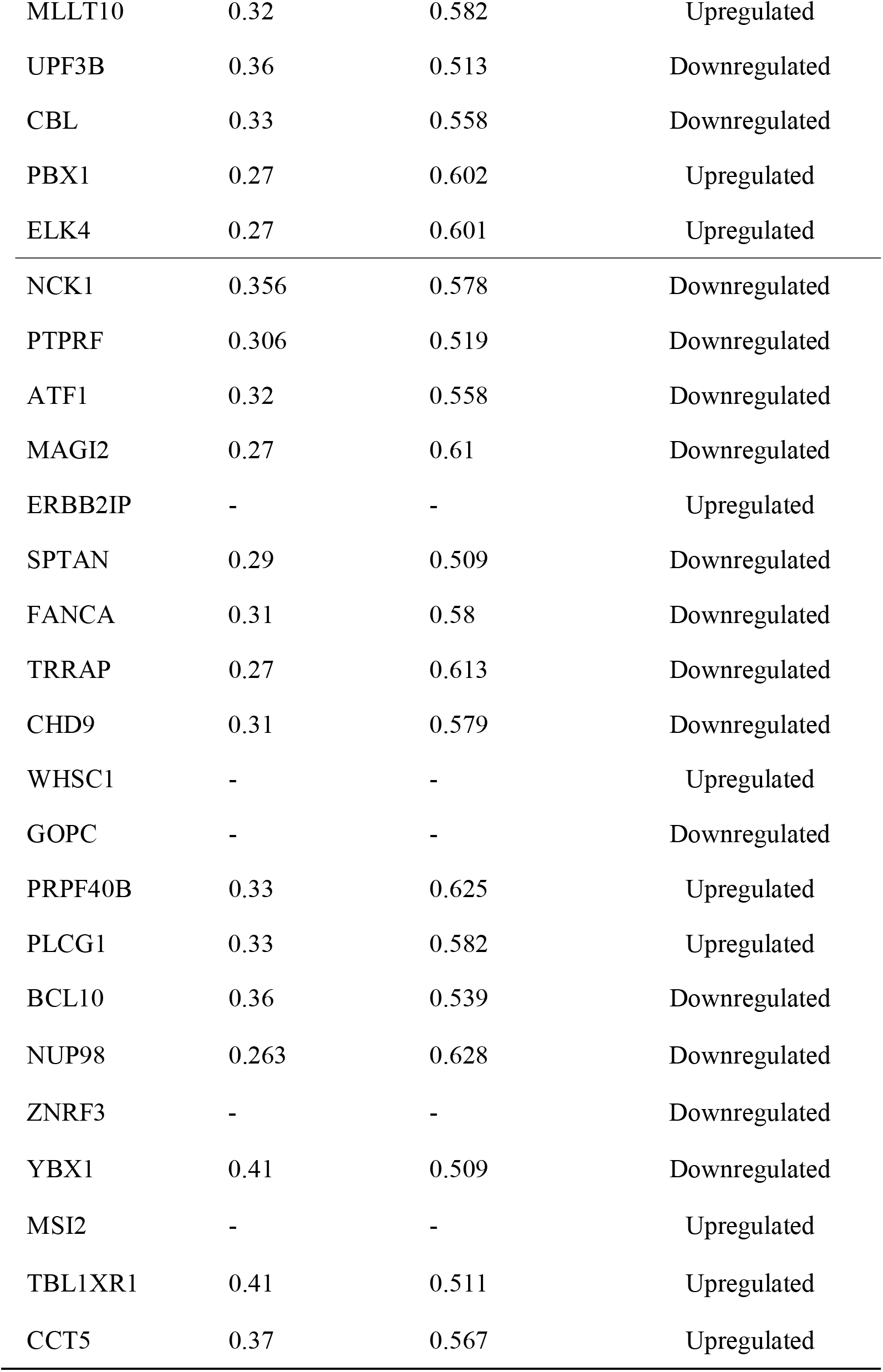
Threshold value between early-late segregation for genes selected by the models

| Training set | Gene | Threshold | ROC | Differential expression |
| --- | --- | --- | --- | --- |
| Selected Gene expression | CDKN1A | 0.32 | 0.56 | Upregulated |
|  | FKBP4 | 0.26 | 0.509 | Upregulated |
|  | DAZ3 | 0.36 | 0.556 | Upregulated |
|  | DNAJA1 | 0.28 | 0.501 | Downregulated |
|  | ECH1 | 0.28 | 0.576 | Upregulated |
|  | RBMY1B | 0.35 | 0.515 | Upregulated |
|  | GABARAP | 0.32 | 0.515 | Downregulated |
|  | MAL2 | 0.31 | 0.611 | Upregulated |
|  | EDA2R | 0.31 | 0.579 | Upregulated |
|  | ITPA | 0.34 | 0.53 | Upregulated |
|  | EIF4A1 | 0.3 | 0.629 | Upregulated |
|  | CTTN | 0.27 | 0.523 | Upregulated |
|  | RAB18 | 0.24 | 0.518 | Upregulated |
|  | GPX4 | 0.45 | 0.535 | Upregulated |
|  | EIF6 | 0.3 | 0.569 | Upregulated |
|  | TIMP1 | 0.34 | 0.572 | Downregulated |
|  | HLA-DQB1 | 0.33 | 0.566 | Downregulated |
|  | TYMP | 0.345 | 0.571 | Upregulated |
|  | PAM | 0.28 | 0.517 | Downregulated |
|  | PGRMC1 | - | - | Downregulated |
| Driver gene expression | DNAJB1 | 0.29 | 0.519 | Upregulated |
|  | GAS7 | - | - | Downregulated |
|  | BCOR | 0.34 | 0.636 | Upregulated |
|  | SKI | 0.25 | 0.573 | Downregulated |
|  | ETV4 | 0.257 | 0.556 | Downregulated |

|  |  |  |  |
| --- | --- | --- | --- |
| MLLT10 | 0.32 | 0.582 | Upregulated |
| UPF3B | 0.36 | 0.513 | Downregulated |
| CBL | 0.33 | 0.558 | Downregulated |
| PBX1 | 0.27 | 0.602 | Upregulated |
| ELK4 | 0.27 | 0.601 | Upregulated |
| NCK1 | 0.356 | 0.578 | Downregulated |
| PTPRF | 0.306 | 0.519 | Downregulated |
| ATF1 | 0.32 | 0.558 | Downregulated |
| MAGI2 | 0.27 | 0.61 | Downregulated |
| ERBB2IP | - | - | Upregulated |
| SPTAN | 0.29 | 0.509 | Downregulated |
| FANCA | 0.31 | 0.58 | Downregulated |
| TRRAP | 0.27 | 0.613 | Downregulated |
| CHD9 | 0.31 | 0.579 | Downregulated |
| WHSC1 | - | - | Upregulated |
| GOPC | - | - | Downregulated |
| PRPF40B | 0.33 | 0.625 | Upregulated |
| PLCG1 | 0.33 | 0.582 | Upregulated |
| BCL10 | 0.36 | 0.539 | Downregulated |
| NUP98 | 0.263 | 0.628 | Downregulated |
| ZNRF3 | - | - | Downregulated |
| YBX1 | 0.41 | 0.509 | Downregulated |
| MSI2 | - | - | Upregulated |
| TBL1XR1 | 0.41 | 0.511 | Upregulated |
| CCT5 | 0.37 | 0.567 | Upregulated |

### Gene Ontology

Clusterprofiler R package was used for gene ontology enrichment analysis of the gene set selected for the complete gene expression-based model and selected gene set for the driver gene-based model. It reveals enrichment in molecular functions such as transferase and hydrolase activity for gene set for the driver gene-based model (Figure 7a). Cathepsin D is a lysosomal hydrolase which is having increased expression in tumors that results in degradation of extracellular matrix causing metastasis [42]. Increased expression of glycoprotein-sialytransferase is associated with altered membrane synthesis resulting in invasiveness and neoplastic state [43].

**Figure 7:**
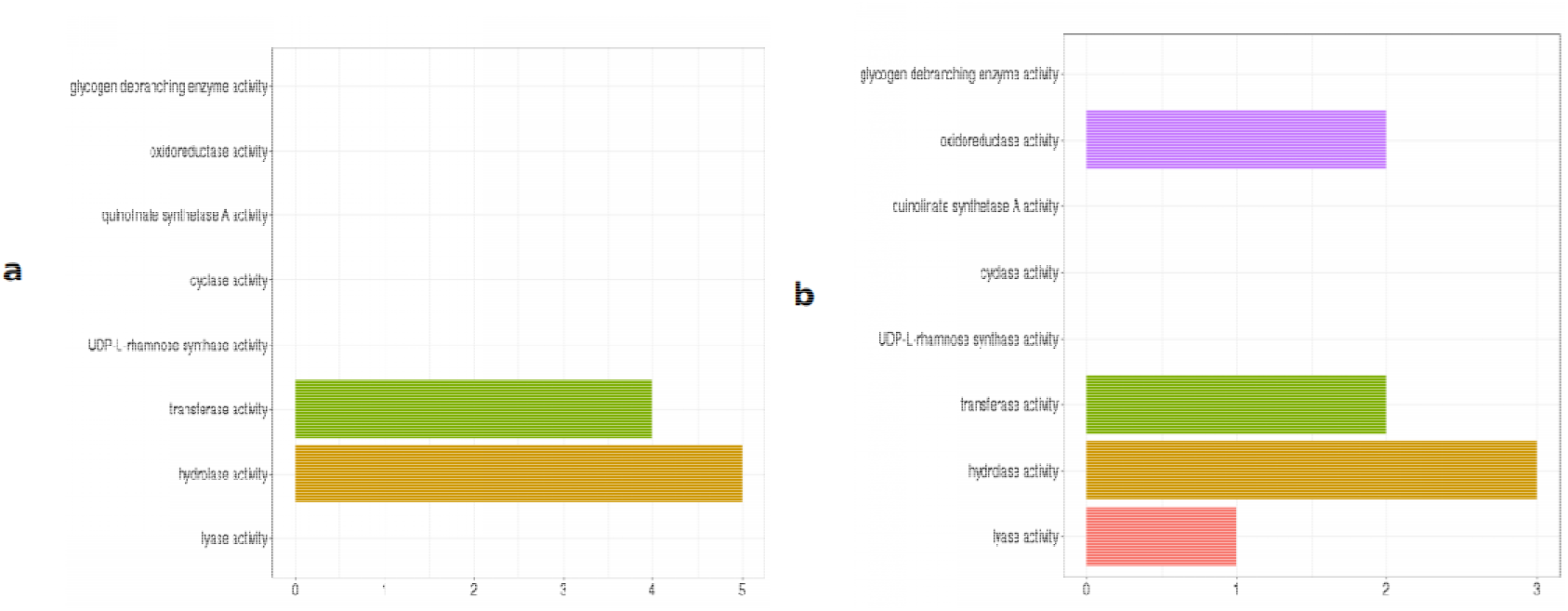
Gene ontology analysis of genes selected for model building. a. Gene ontology analysis of the gene set from driver gene expression-based model for molecular function. b. Gene ontology analysis of gene set from complete gene expression-based model for molecular function.

The selected training gene set for the complete gene expression-based model was found to be enriched in molecular functions related to oxidoreductase activity, lyase, hydrolase and transferase activity (Figure 7b). Glutathione-dependent oxidoreductase-CLIC3 is secreted by cancer cell which contributes to tumour micro-environment by promoting angiogenesis and tumour cell invasion [44]. CSE (Cystathion-gamma-lyase) regulates STAT3 signalling which promotes cell proliferation in breast cancer [45].

The selected training gene set for the complete gene expression-based model was found to be enriched in cellular components related to plasma membrane, endoplasmic-reticulum membrane, organelle membrane and nuclear-endoplasmic reticulum membrane (Figure 8a). Mitochondria-associated ER-membrane responds to various stress signals including apoptotic signalling, inflammatory signalling and unfolded protein response (UPR). These pathways may be perturbed due to abnormal or uncontrolled expression of related genes resulting in cancer development [46]. Training Gene set for the driver gene expression-based model is enriched in cellular component such as plasma, membrane and organelle membrane (Figure 8b).

**Figure 8:**
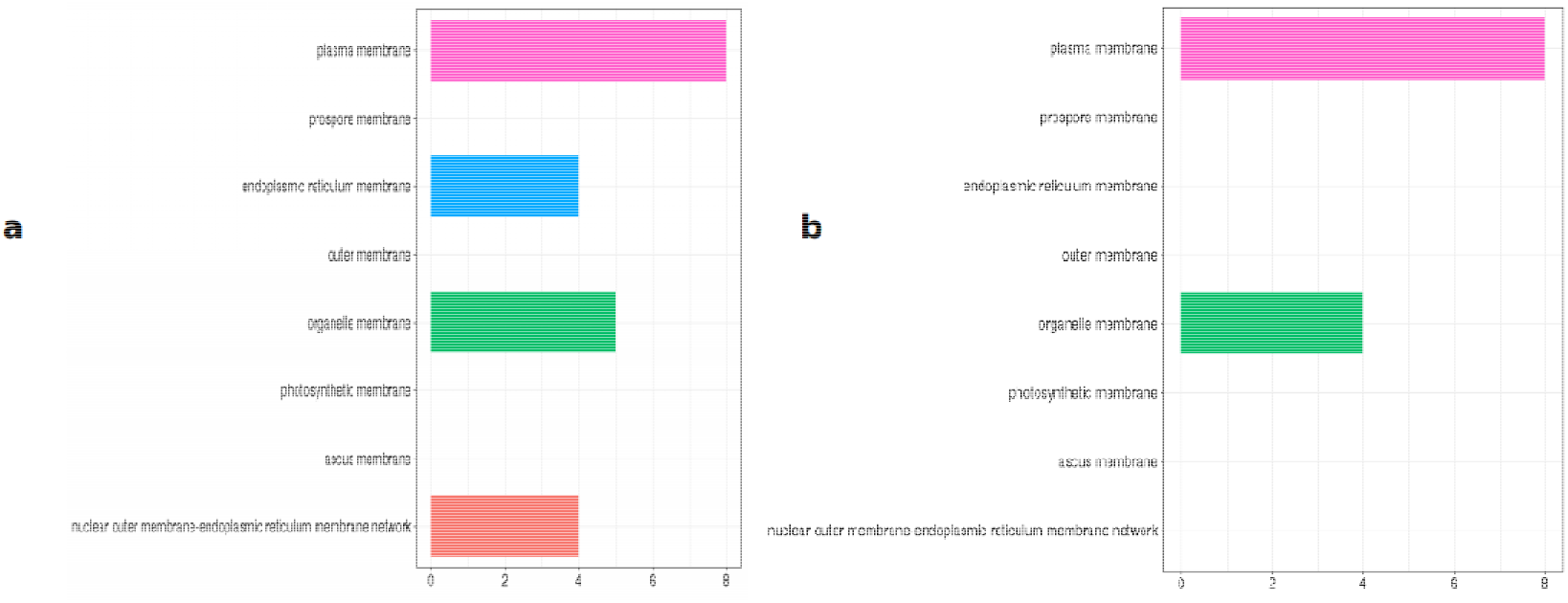
Gene ontology analysis of genes selected for model building. a. Gene ontology analysis of gene set from feature-selection based model for cellular component. b. Gene ontology analysis of gene set by complete gene expression-based model for cellular component.

Gene set from driver gene-based model is enriched in biological processes related to transcriptional misregulation and ErbB signaling (Figure 9a). Transcription factors are involved in tumorigenesis by altering expression profiles of their targets [47]. ErbB tyrosine kinase receptors are found to be activated by epidermal growth factor controlling cellular proliferation, angiogenesis and metastasis in breast cancer [48]. Gene features from the complete Gene expression-based models are more enriched in biological process related to immunological response such as T cell costimulation, immunoglobin response (Figure 9b). Impaired expression of HLA-DQB1 due to change in methylation pattern of gene is associated with esophageal squamous cell carcinoma by altering immune response pattern [49].

**Figure 9:**
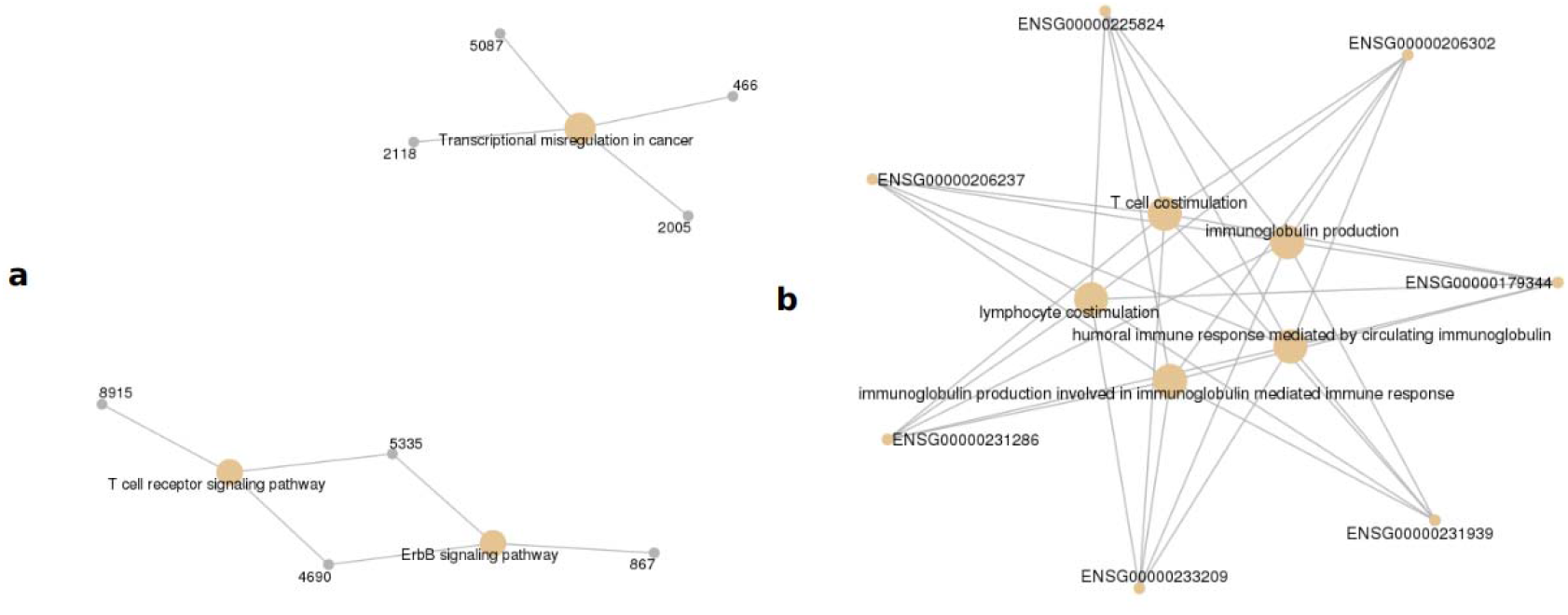
Gene ontology analysis of genes selected for model building. a. Gene ontology analysis of gene set from driver gene expression-based model for biological process. b. Gene ontology analysis of gene set by complete gene expression-based model for biological process.

## METHODS

### Data mining

The study dataset was obtained from TCGA using TCGA2STAT R package, which automatically downloads and processes TCGA genomics and clinical data into a format convenient for statistical analyses in R environment [24]. The package imports and processes molecular profile from high-throughput experiments such as microarray, next generation sequencing and methylation array.

### Data Pre-processing and normalization

As an initial step of pre-processing, which aids in preliminary feature reduction for a feature-rich training dataset, gene features showing near zero variance across the two classes were removed. Near zero variance features are the feature which either have unique value or have few unique value relative to the number of samples. Along similar lines, the features having more than 80% correlation with each other can prove to be problematic for machine-learning. Hence, such feature pairs/groups were also removed in a way where only a single feature of the group remains. These two tasks were performed using the Caret, an R package [25]. We used RPKM (Reads Per Kilobase of transcript per Million mapped reads) values of the reads for supervised machine-learning analysis. RPKM is a measure of normalization of RNA-seq data with the total read length and number of sequencing reads for a given sample [50]. The training datasets for standardized using z-score normalization. It converts all the features to common scale with mean zero and standard deviation 1. The normalized data-set was used for training models.

### Feature selection

Feature selection is an advantageous step before machine-learning which reduces the dimensionality of datasets [23]. Given the possibly large sets of features, it helps in searching for the subset of features that has relevance in terms of a given predictor variable [51]. For example, feature selection has been used to develop a prediction model for prognosis of metastasis and responses to adjuvant therapy in breast cancer, using 50-gene signature [52]. It also helps in improving the accuracy of a classifier by removing irrelevant data [53]. The main challenge associated with current data mining technologies is the high dimensionality of datasets combined with homogenous nature of data [54].

For reducing the dimensionality of the datasets and identifying relevant features for building efficient machine-learning classifiers, we implemented various feature selection algorithms such as RFE, RLASSO, random forest, linear modelling and linear regression, which provide individual ranking to gene features. Recursive Feature Extraction (RFE) is a method that utilizes recursion for feature extraction where smaller and smaller sets are considered as features until the desired number of features is returned. Randomized lasso is a stability selection method, which is combination of sub-sampling of high dimensional datasets and selection algorithm [55]. Linear regression assumes that features that are important have highest coefficient in the model, and features that have low importance have lower coefficient in the model. When there are multiple correlated features, small change in data can lead to large change in model. Regression model uses regularization method that adds an additional penalty to a model in order to minimize the sum of squared error of training model using lasso and ridge regression methods. Lasso regression methods performs L1 regularization minimizes absolute sum of the coefficient and producing sparse solution. Ridge regression performs L2 regularization minimizing squared absolute sum of the coefficients. The Least absolute shrinkage and selection operator (LASSO) does regression analysis for parameter estimation and variable selection simultaneously [56]. Random forest uses decision tree based strategies to rank feature based on attribute “feature importance”. There are reports of use of Random forest algorithm to extract relevant feature from biological datasets such as gene expression, methylation [57]. In this work, the feature selection methods were implemented using the Python 3.6 scikit-learn library.

These feature-selection methods were used to rank the gene features of the training datasets. All the methods were implemented using the popular Python 3.6 scikit-learn library. All of the above-mentioned methods report individual ranking for the features. In order to get consensus ranking, we calculated the overall mean of each feature rank obtained from individual method. Subsequently, the top 50, 60, 80 and 100 features were used to train and evaluate accuracy of models for binary classification of early versus late IDC, based on 5 machine-learning methods namely - RF, Naive Bayes, SVM, Logistic regression (LR) and Decision tree. Gene features list that gave the highest accuracy for all the machine-learning method, were selected for model generation and evaluation. t-SNE technique was used for visualization of our gene expression datasets returned after feature selection to check if datasets are segregating into defined class based on selected features for visualization of high dimensional data-point t-SNE uses random walk on neighbourhood graph that allows implicit structure of data point to influence the way groups of data is present [58].

### Handling data imbalance

Real world datasets have higher composition of ‘normal class’ as compared to ‘abnormal class’, introducing bias in classification model. Combination of over-sampling of minority class along with under-sampling of majority class can aid in increasing the classifier performance. To check the SMOTE resampling, models were trained on datasets where SMOTE resampling was employed.

### Training classification models

We have used Supervised machine learning algorithm in this work which targets to classify class label by building decision rule [59]. After feature selection and data processing, we trained different algorithms to generate efficient classifiers for early and late tumour stage. We used five different algorithms – Random Forest, Naive Bayes, LinSVM (Support Vector Machines with linear kernels), Logistic regression and Decision tree. Naive Bayes is based on Bayesian theorem that calculates the probability of attribute to fall in particular instance with the assumption that every attribute is independent from other attributes [60]. Random forest uses ensemble of decision tree by random selection of features to split node [61]. Random forest performs sub-sampling of training datasets to construct large number of trees for solving regression problem [62]. SVM implements Sequential Optimization Algorithm for decision function [63].

#### Training-cum-validation

The five supervised machine-learning algorithms (Random Forest, Naive Bayes, LinSVM, Logistic regression and Decision tree) were trained on subset of features obtained from feature selection and validated by 10-fold cross validation. The training models were compared by their accuracy, auROC, precision -recall and F-measure value.

#### Independent data testing

We further re-evaluated the best-trained model on an independent dataset which was not used in the classifier training at all.

### Calculating threshold expression values for selected gene features

We performed differential expression analysis for the selected gene features by the two models, for early-late datasets to find out the differential expression of gene features selected by our model. Each gene feature selected by our model had range of expression across all the samples. We executed machine-learning and model evaluation for every single feature selected by our classifiers with threshold set across its expression range. The value that was giving highest ROC was considered as threshold value of expression value that could discriminate between early-late stages. Threshold value is the expression value beyond which the sample will segregate into two groups, in our case ‘early’ and ‘late’ stage.

### Cancer driver gene expression-based model

The available driver gene list for the cancer were also used for building model to discriminate early-late stages of breast cancer. We complied list of driver genes using Cosmic, intoGen and baile, which is expert curated list of driver genes in human cancers. Cosmic stands for catalogue of somatic mutations in cancer which is expert curated list of driver gene in human cancer which is widely used in medical research [26]. IntoGen identifies somatic mutation, gene, pathway that are involved in tumorigenesis by analysis of 13 cancer. [27] Bailey list identifies 299 molecular cancer gene by pan-cancer and pan-software analysis of 9,423 tumour exome using 26 computational tool [28]. We reduced the data-set to these gene features, which was then used for feature selection and model building repeating the abovementioned steps to generate driver gene expression-based model for web server.

### Gene Ontology

GO was performed on the list of genes returned by the feature selection methods to determine which gene families play role in the progression of breast cancer. We performed enrichment analysis using clusterprofiler R package. The package makes use of the datasets from the post genomic era high throughput technologies such as RNA-seq, micro-array, etc. to examine cellular molecules at systems level [64]. We also performed string protein-protein interaction analysis to discover major pathways targeted by selected gene features.

### External data-set evaluation

To further check the performance of our model, we obtained independent datasets form GEO with accession ID GSE61304 containing 60 samples of ductal carcinoma with clinical stage information obtained using microarray profiling. GEOquery package helps the user to access the information stored in GEO directly using Bioconductor without any formatting or parsing problem [65]. Biomart was used to annotate the probe IDs of microarray datasets with gene symbol [66]. If a particular probe is sequenced multiple times, WGCNA R package collapserow function which uses bio-statistical methods to select one single representative row of each probe id [67]. Subsequently, RMA normalization was performed using GRCMA R package converting the expression in log 2 scale to make its distribution comparable to RNA-seq datasets [68,69]. This independent-testing dataset were segregated into driver-testing datasets and feature-testing datasets for performance evaluation of the generated models were evaluated.

## Conclusion

We have successfully applied supervised machine-learning classification on gene expression profiles to develop classification models for discrimination between early and late stage of invasive ductal carcinoma. The RNA-seq data obtained from TCGA had various information related to samples from age, survivability, TNM staging, histological subtype and pathological stage in the form of metadata or clinical data.

The data yielded 20,505 gene expression used as training features to be considered for classification model trainings. This voluminous dimensionality was facilitated using various data pre-processing and feature selection methods. After this, the classifier models were generated by applying various machine-learning algorithms. Based on trained classifiers, we developed a web-server Duct-BRCA-CSP which predicts the inputs sample to be in early or late stages using selected gene expression profile in a sample. The model trained on gene features shortlisted by the feature selection methods can reliably differentiate samples between early and late stage with high accuracy. The shortlisted genes also validates candidate biomarkers and potential biomarkers for development of improved diagnosis, prognosis and treatment of IDC patients. The combined power of machine-learning and next generation sequencing can also provide important insights into the progression of breast cancer from early to late stage.

## Discussion

We developed a web-server Duct-BRCA-CSP for invasive ductal carcinoma which predicts tumour stage of a sample on the basis of RNA-seq expression profile, rather than its tumour size, imaging or survivability. Our study is preliminary in nature, however, in the future, the availability of datasets from higher number of patients, especially those representing late stage may help in building more efficient stage specific. In addition, further inclusion of additional datasets such as mutation profile, methylation data and protein isoform data may also improve the accuracy of classifiers. Inclusion of paired datasets can also further aid in gaining insights into the progression of breast cancer. To the best of our knowledge, the webserver Duct-BRCA-CSP is a server which is first of its kind for prediction of IDC tumour stages based on gene expression profile.

## Supporting information

Supplementary file III

## Supplementary information

### Supplementary file I

This file consists of four figures – Figure S1 is a distribution plot of gene feature DNAJB1 Before normalization, Figure S2 is a distribution plot of gene feature DNAJB1 after normalization, Figure S3 is Heatmap of differential expression between early and late IDC stages for the gene set from complete gene expression-based model, Figure S4 is Heatmap of differential expression between early and late IDC stages for the gene set from driver gene expression-based model. Figure S5: Due to high class imbalance (461 early stage versus 161 late stage) Synthetic Minority Oversampling technique (SMOTE) was employed used python scikit-learn. Scatter plot to evaluate effectiveness of SMOTE + ENN re-sampling technique to handle class imbalance. Early stage datasets are labelled as #1, and late stage labelled as #0. a) Prior to resampling b.) Post SMOTE resampling. As compared to original sample prior to resampling, SMOTE resampling rendered a larger late stage sample using k-neighbour of majority class. Figure S6: SMOTE resampled datasets were used to train the binary classification model and their accuracy was again evaluated. Most of the algorithms displayed improved accuracy of classification after SMOTE resampling. Accuracy of machine learning algorithm before SMOTE resampling and after SMOTE resampling is shown here. **NB:** Naïve Bayes, **LR:** Logistic Regression, **RF:** Random Forest, **SVM:** Support Vector Machine, **DT:** Decision Tree. **BS:** Before SMOTE resampling **AS:** After SMOTE resampling. **X axis:** Machine learning algorithm, **Y axis:** Accuracy.

### Supplementary file II

Analysis of selected gene feature for each model for Overall Survival Kaplan-Meier Estimate from cBioportal for Cancer Genomics. The file consists of two figures – Figure S7: Survival Kaplan-Meier estimate for gene set from driver gene expression-based model. Figure S8: Survival Kaplan-Meier estimate for gene set from complete gene expression-based model. This file is in a Microsoft word format.

### Supplementary file III

Gene symbols and literature validation of selected genes. The Microsft word formatted file consists of 50 genes selected for generation of training dataset for gene expression-based model and driver gene expression-based model.

## Author Contribution

Conceptualization, S.R. D.G.; Methodology, V.G.; Software, R.K.; Validation, S.R., V.G. and R.K.; Formal Analysis, S.R.; Investigation, S.R, R.K.; Resources, S.R.; Data Curation, S.R.; Writing – Original Draft Preparation, S.R.; Writing – Review & Editing, S.R, D.G; Visualization, S.R.; Supervision, D.G.; Project Administration, D.G; Funding Acquisition, D.G.

## Acknowledgements

This work was financially supported by the Department of Biotechnology (DBT), Government of India, grants BT/PR6963/BID/7/427/2012 and BT/BI/25/066/2012 awarded to DG.

