## Supplementary file I for "Classification models for Invasive Ductal Carcinoma Progression, based on gene expression data-trained supervised machine learning"

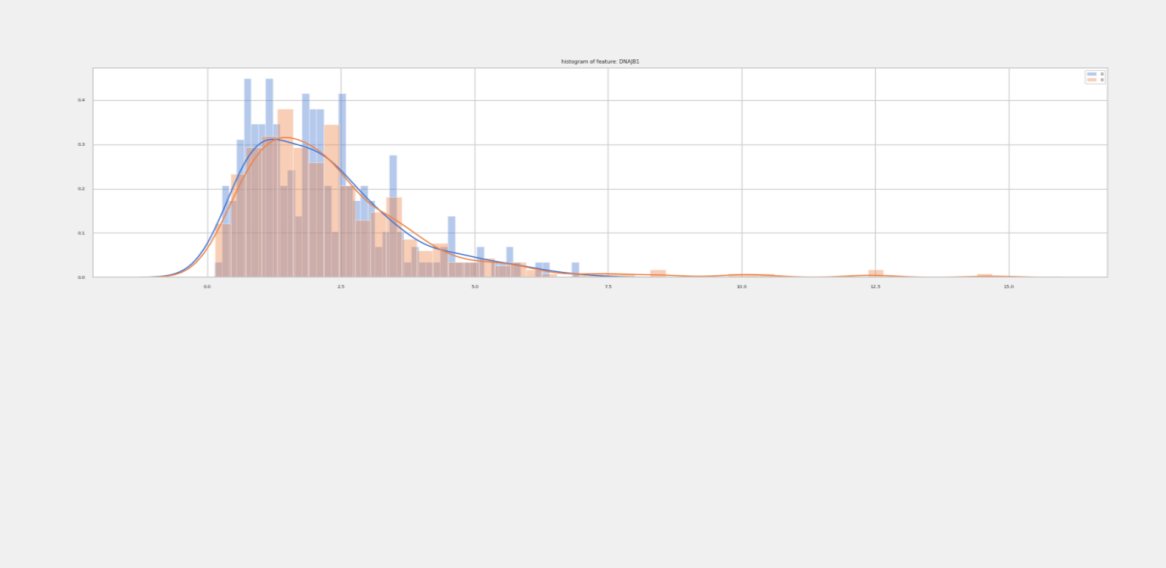

Figure S1: Distribution plot of gene DNAJB1 expression before normalization

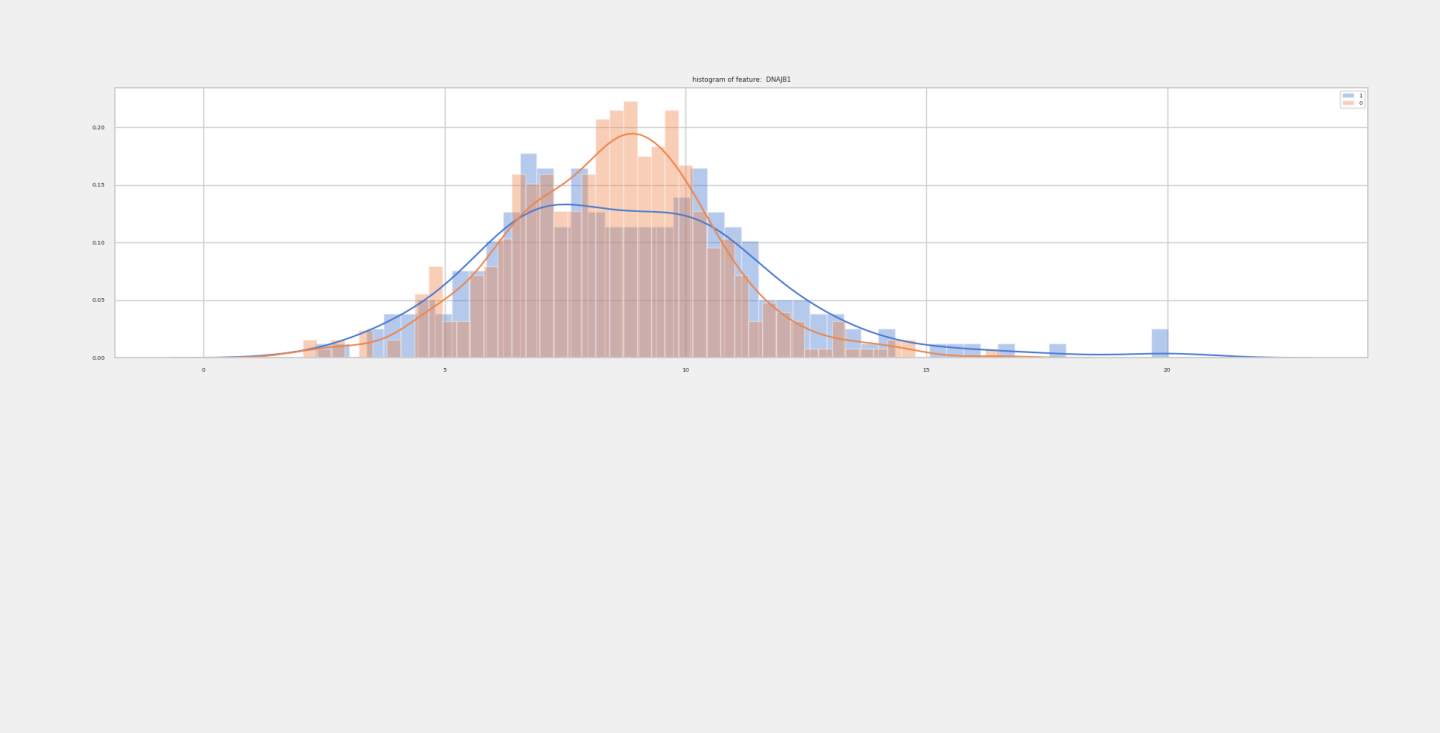

Figure S2: Distribution plot of gene DNAJB1 expression after normalization

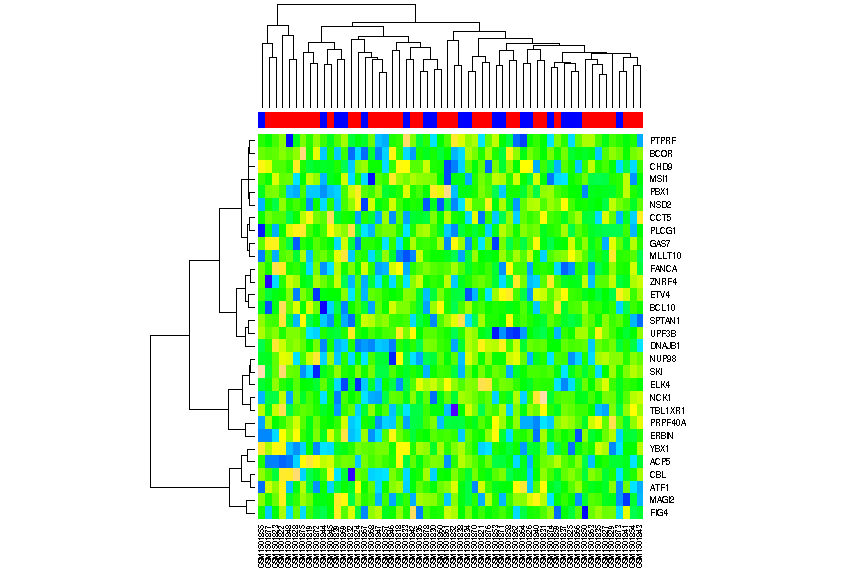

Figure S3: Heatmap of differential expression between early and late IDC stages for the genes set from complete gene expression-based model.

Red: Early stage Blue: Late stage

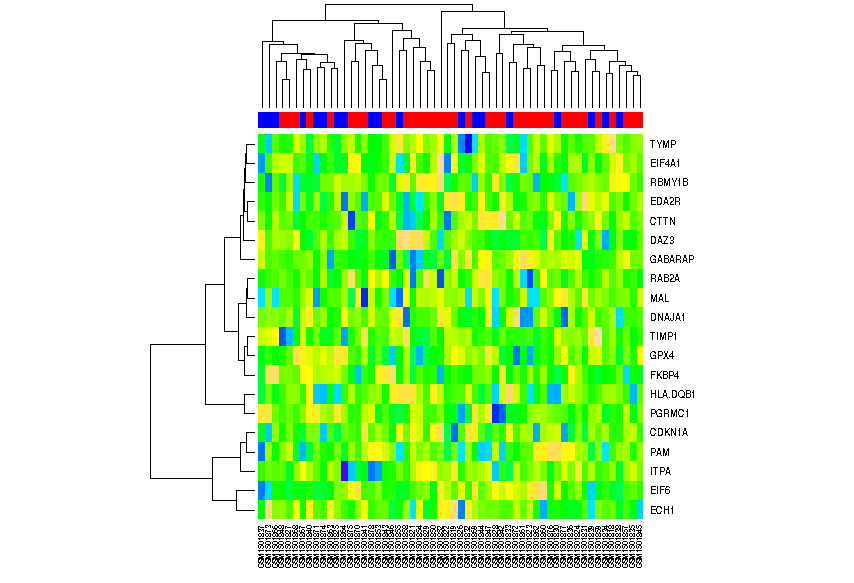

Figure S4: Heatmap of differential expression between early and late IDC stages for gene set from driver gene expression-based model.

Red: Early stage Blue: Late stage

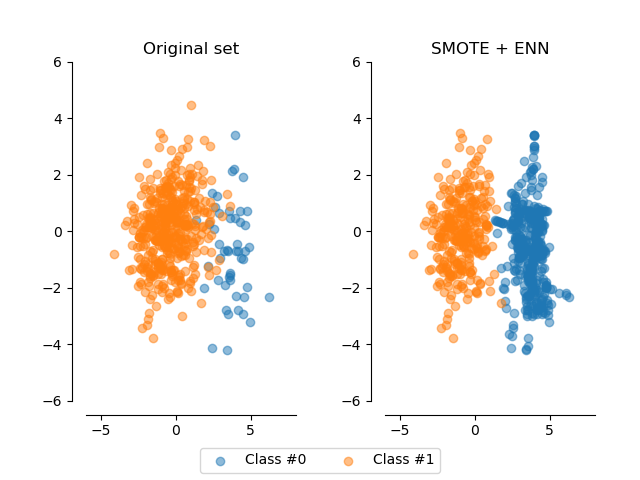

Figure S5: Due to high class imbalance (461 early stage versus 161 late stage) Synthetic Minority Oversampling technique (SMOTE) was employed used python scikit-learn. Scatter plot to evaluate effectiveness of SMOTE + ENN re-sampling technique to handle class imbalance. Early stage datasets are labelled as #1, and late stage labelled as #0. a) Prior to resampling b.) Post SMOTE resampling. As compared to original sample prior to resampling, post SMOTE resampling the more late stage samples are generated using k- neighbour of majority class.

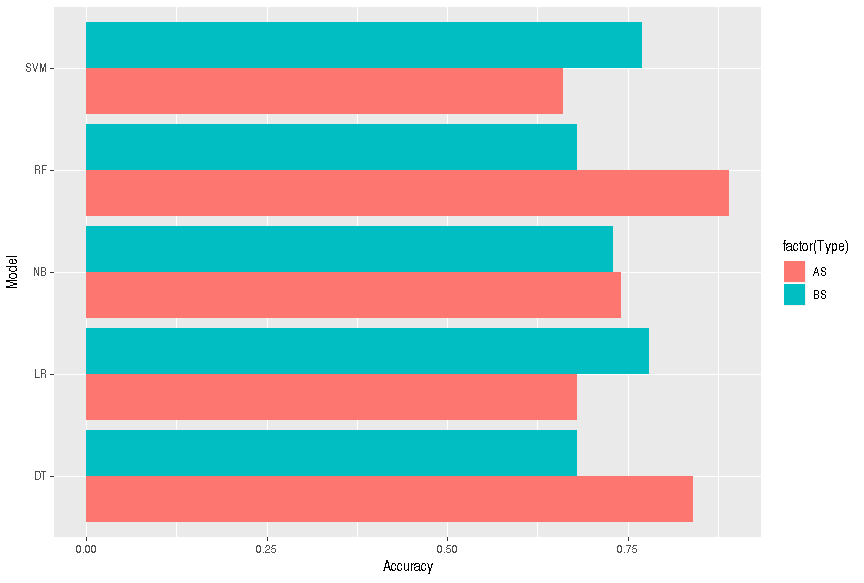

Figure S6: SMOTE resampled datasets were used to train the binary classification model and their accuracy was again evaluated. Majority of algorithm was showing improved accuracy of classification model after SMOTE resampling. Accuracy of machine learning algorithm before SMOTE resampling and after SMOTE resampling. NB: Naïve Bayes, LR: Logistic Regression, RF: Random Forest, SVM: Support Vector Machine, DT: Decision Tree.BS: Before SMOTE resampling AS: After SMOTE resampling. X axis: Machine learning algorithm Y axis: Accuracy.
