## Supplementary file III for "Classification models for Invasive Ductal Carcinoma Progression, based on gene expression data-trained supervised machine learning"

| S.No. | Gene Symbol | Associated Cancer Type | Remarks and References |
| --- | --- | --- | --- |
| 1. | DNAJB1 | Breast carcinoma, hepatocellular carcinoma | It negatively regulates MIG6 which is tumor suppressor that results in cancer cell proliferation by EGFR signaling pathway.[1] |
| 2. | GAS7 | Breast cancer | It is directly regulated by p53 and suppresses metastasis in breast cancer.[2] |
| 3. | BCOR | T-cell lymphoblastic leukemia | It function as tumor suppressor and its mutation has been identified to be associated with various hematological malignancies.[3] |
| 4. | SKI | Breast cancer | Its divergent expression is associated with fibroblast at tumor microenvironment in breast cancer.[4] |
| 5. | ETV4 | Breast cancer | Its overexpression is associated with distant metastasis and poor prognosis in breast cancer.[5] |
| 6. | MLLT10 | T- acute lymphoblastic leukemia | MLLT10 has specific leukemia fusions in T- acute lymphoblastic leukemia.[6] |
| 7. | UPF3B | Different cancers | Role in nonsense-mediated mRNA decay and cancer.[7] |
| 8. | CBL | Breast cancer | It functions as an adaptor protein with E3 ubiquitin ligase and mutation is associated with carcinogenesis.[8] |
| 9. | PBX1 | Breast Cancer | Its expression is associated with ER meditated transcriptional response in driving breast cancer.[9] |
| 10. | ELK4 | Breast Cancer | It is ETS transcription factor whose copy number changes, associated with pathogenesis of cancer.[10] |
| 11. | NCK1 | Breast cancer,  Lung cancer | It regulates breast cancer prognosis and metastasis by growth and vascularization of primary tumor.[11] |
| 12. | PTPRF | Different cancers | Important cell cycle regulator associated with tyrosine kinase and have role in cell proliferation, apoptosis, migration and invasion.[12] |
| 13. | ATF1 | Breast cancer | Transcription factor that regulates cell cycle and apoptosis and is involved in cancer progression.[13] |
| 14. | MAGI2 | Breast cancer | It serves as tumor suppressor and has decreased expression in breast cancer.[14] |
| 15. | ERBB2IP | Breast cancer | It belongs to family of receptor tyrosine kinase which regulates breast tumor formation and progression.[15] |
| 16. | SPTAN1 | Gastric cancer | Its different expression is linked to gastric cancers.[16] |
| 17. | FANCA | Breast cancer | FANCA gene duplication is associated with increased risk of breast cancer.[17] |
| 18. | TRRAP | Ovarian cancer | Its over-expression is associated with increased proliferation and stemness of ovarian cancer.[18] |
| 19. | CHD9 | Breast cancer | It belongs to family of chromatin regulator whose inactivation results in distortion to cellular machinery related to transcription and DNA damage response, resulting in inducing cancer.[19] |
| 20. | WHSC1 | Cervical cancer | Its hypomethylation results in its over-expression that promotes cervical carcinogenesis.[20] |
| 21. | GOPC | Colorectal cancer | Its under-expression is associated with venous invasion and poor prognosis in colorectal cancer.[21] |
| 22. | PRPF40B | Acute myeloid leukemia | It is a spliceosome complex gene whose mutation is associated with acute myeloid leukemia.[22] |
| 23. | NUP98 | Murine carcinoma, Hepatocellular carcinoma | It regulates the expression of p21 and has reduced expression in hepatocellular carcinoma.[23] |
| 24. | CCT5 | Breast cancer | It has increased expression in p53 mutated breast cancer and also implicates resistance to docetaxel treatment in breast cancer.[24] |
| 25. | PLCG1 | Cancer | It is phospholipase enzyme that promotes cell invasion, metastasis and tumor progression in cancer.[25] |
| 26. | BCL10 | Ovarian cancer | It has role in DNA damage response and associated with tumor aggression and poorer prognosis.[26] |
| 27. | ZNRF3 | Osteosarcoma | The gene is targeted by miRNAs. [27] |
| 28. | YBX1 | Breast Cancer | It is transcription and translation related protein found to overexpressed in various malignancies.[28] |
| 29. | MSI2 | Breast cancer | It is upstream regulator of ESR1 that modulated estrogen receptor pathway resulting in cancer cell growth.[29] |
| 30. | TBL1XR1 | Gastric cancer | It is found to be overexpressed in gastric cancer and can serve as therapeutic target.[30] |
| 31. | CDKN1A | Breast cancer | Its over-expression correlates significantly with tumor size and lymph node metastasis in breast cancer.[31] |
| 32. | FKBP4 | Prostate cancer | The protein enhances the transcriptional activity of androgen receptor signaling, overexpressed in CRPC.[32] |
| 33. | DAZ3 | Testicular germ cell tumor | Its copy number changes is associated with testicular germ cell tumor.[33] |
| 34. | DNAJA1 | Breast cancer | It controls mutant p53 whose stabilization contribute to malignancy in breast cancer.[34] |
| 35. | ECH1 | Breast cancer | Its increased expression results in PPAR increased expression promotes adipogenesis in breast cancer cell.[35] |
| 36. | RBMY1B | liver cancer | It is member of RNA binding protein that is having male specific activation in liver cancer.[36] |
| 37. | GABARAP | Breast Cancer | It is involved in receptor transport and autophagy and associated with lymph node metastasis in breast cancer.[37] |
| 38. | MAL2 | Ovarian cancer, Cervical cancer | Its down-regulation results in redistribution of lipid raft in cancer. [38] |
| 39. | EDA2R | Breast cancer | Its transcriptional activation results in cell death in breast cancer.[39] |
| 40. | ITPA | Different cancers | It encodes enzyme that is responsible for deamination of adenine resulting in mutagenesis and DNA damage response.[40] |
| 41. | EIF4A1 | Breast cancer | It encodes protein that is involved in initiation of protein synthesis and its dys-regulation results in tumorigenesis.[41] |
| 42. | CTTN | Melonoma, Breast cancer, Colorectal cancer | It encodes cortactin, which is a regulator of actin cytoskeletal whose amplification results in metastatic breast cancer. [42] |
| 43. | GPX4 | Clear cell carcioma | It serves as therapeutic target for clear-cell carcinoma.[43] |
| 44. | RAB18 | Breast cancer | It is regulated by miRNA whose dys-regulation results breast cancer proliferation and invasion.[44] |
| 45. | EIF6 | Colorectal cancer,  Head and neck  carcinoma | Its overexpression results in activation of Wnt signaling pathway which dys-regulates cell division and promotes oncogenesis.[45] |
| 46. | TIMP1 | Breast cancer | It encodes protein that directly regulates apoptosis and metastasis in triple negative breast cancer.[46] |
| 47. | HLA-DQB1 | Breast cancer | Its allele is found to have protective role in immune surveillance process of breast cancer.[47] |
| 48. | TYMP | Rectal cancer | It is over-expressed in cancer and promotes process related to angiogenesis.[48] |
| 49. | PAM | Lung cancer, prostate cancer | Increased activity in Lung and Prostate cancers.[49] |
| 50. | PGRMC1 | Breast cancer, Ovarian cancer | It is found to have mutually exclusive expression with ER in breast cancer.[50] |
