## Supplementary file II for "Classification models for Invasive Ductal Carcinoma Progression, based on gene expression data-trained supervised machine learning"

***Survival curve for each model***

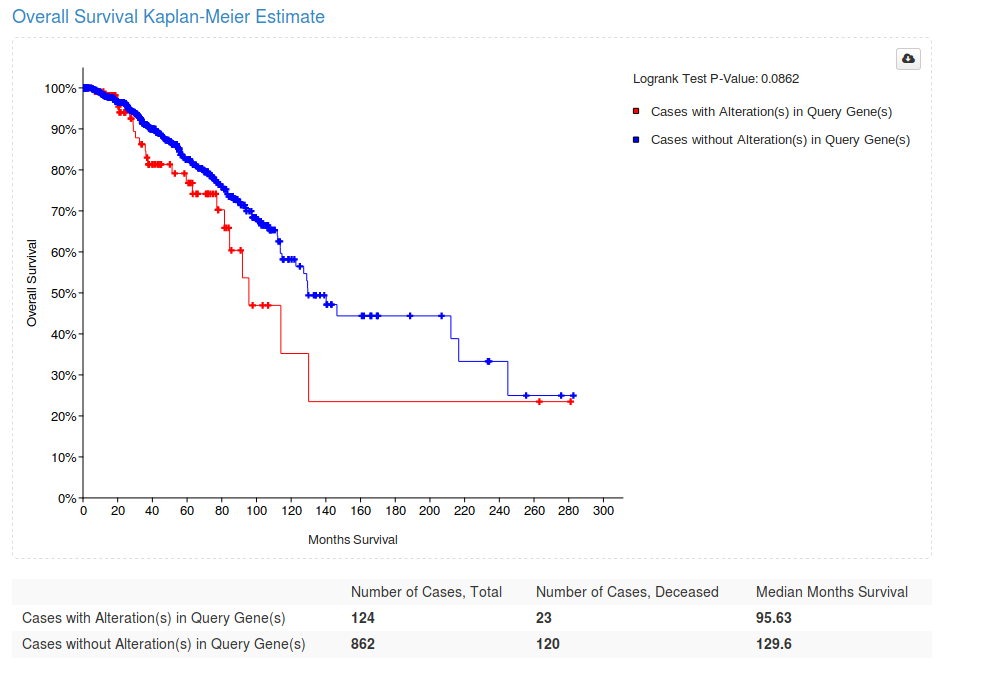

Figure S7: Survival curve for gene set from driver gene expression-based model

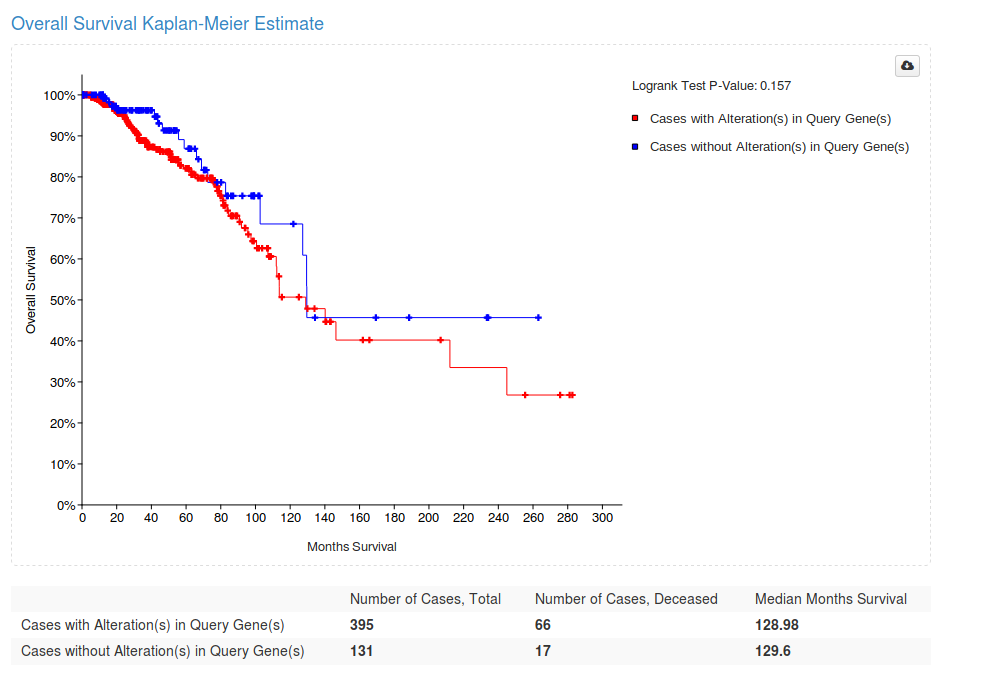

Figure S8: Survival curve for gene set from complete gene-expression based gene model.
